# PANK2-mediated *de novo* CoA synthesis is required for metabolic switching to fatty acid oxidation

**DOI:** 10.1101/2025.10.07.680981

**Authors:** Sandra M. Nordlie, Robert Schafer, Adam Rauckhorst, Ana M. Santos-Exposito, Andres Prieto-Rodriguez, Bryce A Wilson, Idil A. Evans, Una Hadziamehtovic, Frances Nowlen, Carlos Santos-Ocaña, Eric B. Taylor, Michael C Kruer, Sergio Padilla-Lopez

## Abstract

Humans have 3 different PANK enzymes (PANK1-3) that catalyze the first step in the *de novo* synthesis of Coenzyme A (CoA). All PANKs are feedback inhibited by acyl-CoAs but only PANK2 can overcome this inhibition by binding palmitoyl-carnitine. Previous studies, conducted under glucose-replete conditions, have failed to detect a PANK2-mediated contribution to CoA synthesis. We found that exposure to BSA-conjugated palmitate (PAL-BSA) led to activation of fatty acid oxidation (FAO) and the accumulation of both palmitoyl-carnitine and palmitoyl-CoA in HEK293T cells, suggesting that PANK2 is active under these conditions. Isotope tracing experiments with ^13^C_15_N-pantothenate showed that PANK2 uniquely sustains *de novo* CoA synthesis and high production of Acetyl-CoA in the presence of long-chain fatty acids, indicating that FAO is limited by CoA availability in these conditions. Consistent with this mechanism, fibroblasts from PKAN patients exhibited impaired oxidation of palmitoyl-carnitine, confirming the functional relevance of our results in a disease context.

## INTRODUCTION

Coenzyme A (CoA) is an essential cofactor for over a hundred metabolic reactions that span the tricarboxylic acid (TCA) cycle, lipogenesis, lipolysis, β-oxidation and amino acid degradation [1]. CoA’s primary function is to serve as an acyl group carrier via its terminal thiol (-SH) group, enabling the formation of high-energy thioester bonds. These reactions are fundamental to both catabolic and anabolic metabolism, making CoA indispensable for cellular function and systemic homeostasis. CoA synthesis, availability, and regulation are critically relevant for human disease [2]. In cancer, CoA-derived acetyl-CoA fuels lipid biosynthesis [3] and histone acetylation, supporting proliferation and epigenetic remodeling [4]. Enzymes like ATP citrate lyase (ACLY) and acetyl-CoA synthetase 2 (ACSS2) are often upregulated in tumors, making them potential therapeutic targets [5]. In cardiovascular disease, CoA is essential for myocardial fatty acid oxidation and energy production, while its involvement in lipid and cholesterol synthesis links it to atherogenesis [6]. Mutations in CoA biosynthesis enzymes PPCS or PPCDC lead to monogenic forms of dilated cardiomyopathy (MIM #618189) [7]. In diabetes and metabolic syndrome, CoA governs gluconeogenesis, β-oxidation, and insulin sensitivity; dysregulation disrupts metabolic flexibility and contributes to hyperglycemia and insulin resistance [8]. In neurological diseases, CoA is critical for neurotransmitter synthesis and mitochondrial metabolism. Mutations in CoA synthesis enzymes PANK2 and COASY cause the neurodegenerative diseases pantothenate kinase associated neurodegeneration (PKAN; MIM #234200) and COASY protein associated neurodegeneration (CoPAN; MIM #615643) respectively [9-11]. These findings underscore the importance of CoA homeostasis in human health and its potential as a therapeutic target across multiple disease states.

CoA synthesis and degradation are highly regulated in the cell [12] and one of the most important points of control is the phosphorylation of pantothenate (vitamin B5) by pantothenate kinase (PANK) isoforms in the first step of the biosynthetic pathway. The genes for PANKs1-3 yield four catalytically active (kinase) PANK isoforms; PANK1α, PANK1β, PANK2 and PANK3 [13]. PANK1’s α and β isoforms arise from different initiation exons within the *PANK1* gene [14]. PANK1β and PANK3 are 79% identical at the sequence level, but differentially regulated [15]. Localization studies have shown PANK1α within the nucleus, while PANK1β and PANK3 are cytosolic, with a portion of PANK1β associated with clathrin-associated vesicles and recycling endosomes [16]. Interestingly, PANK2 is the most sensitive to feedback inhibition by acetyl-CoA and long-chain Acyl-CoAs such as palmitoyl-CoA, however this inhibition can be reversed by palmitoyl-carnitine [17]. Moreover, PANK2 is the only PANK localized in the mitochondrial intermembrane space, where it is predicted to encounter high concentrations of long chain acyl-carnitines (such as palmitoyl-carnitine) [17]. Finally, a recent study by Dibble and colleagues discovered that PANK4, previously thought to represent a catalytically inactive PANK, actually serves as a phosphatase that depletes the CoA synthesis intermediate 4-phosphopantethine, effectively reducing the production of CoA and indicating a new mechanism to control cellular CoA abundance [18].

Pantothenate kinase associated neurodegeneration (PKAN) is a neurodegenerative disease classified as a form of neurodegeneration with brain iron accumulation (NBIA). Characterized by progressive dystonia and dementia, PKAN features hallmark pathological features that include iron deposition and neuronal loss in the globus pallidus, leading to the characteristic “eye of the tiger” sign on MRI [11]. Although it is well established that biallelic mutations in PANK2 lead to PKAN, the molecular mechanism(s) that lead to PKAN have remained elusive as prior studies have been unable to detect any differences in cellular CoA synthesis or abundance after deletion or overexpression of PANK2 in human cell lines [18-19]. Moreover, studies with mouse PANK2 KO models obtained similar results [20-21]. This has led to uncertainty about whether PKAN is truly caused by a disruption of CoA biosynthesis. Similarly, it has been unclear whether the activity of PANK2 is redundant with the other PANKs and whether PANK2 deficiency could be compensated by PANK1 or PANK3. The answer to these questions has major implications for both our understanding of CoA availability as a regulator of metabolism and the development of rational therapeutic approaches for PKAN. In our studies below, we describe for the first time specific cellular conditions where PANK2 is essential for *de novo* CoA synthesis and outline the cellular consequences of PANK2 deficiency under these conditions.

## RESULTS

### [^13^C_3_ ^15^N_1_]-pantothenate tracing reveals a critical role of PANK2 in de novo CoA synthesis during PAL-BSA exposure

Prior studies indicated that PANK2 can localize in the intermembrane space (IMS) of the mitochondria [16] (Figure 2B), where it is most likely to be exposed to the activating action of Long Chain Acyl-carnitines [17] (Figure 1a), especially when Fatty Acid Oxidation (FAO) is active and production of Acyl-carnitines by the action of carnitine palmitoyl transferase 1A (CPT1A) is high. We sought to create the optimal cellular conditions for PANK2 activity by inducing FAO in HEK293T cells with BSA-conjugated fatty acids as described before [24]. Therefore, we treated HEK293T control cells with BSA-palmitate to induce a metabolic switch from predominant glucose utilization to FAO (Figure 1a). We confirmed that switching had occurred by observing an increase in CPT1A both at the mRNA and protein level along with an increase in acetyl-CoA production (Figure 1a), as activation of fatty acid oxidation normally leads to an increased yield of acetyl-CoA compared to glucose oxidation [6, 19, 25-26]. We confirmed that PAL-BSA incubation led to increased uptake and activation of PAL-BSA to palmitoyl-CoA (Figure 1c) by acyl-CoA synthetases as well as an increase in CPT1A-mediated palmitoyl-carnitine production (Figure 1c) above baseline conditions (BSA vehicle). We anticipated that this increase in palmitoyl-carnitine abundance could stimulate PANK2 activity.

**Figure 1.**
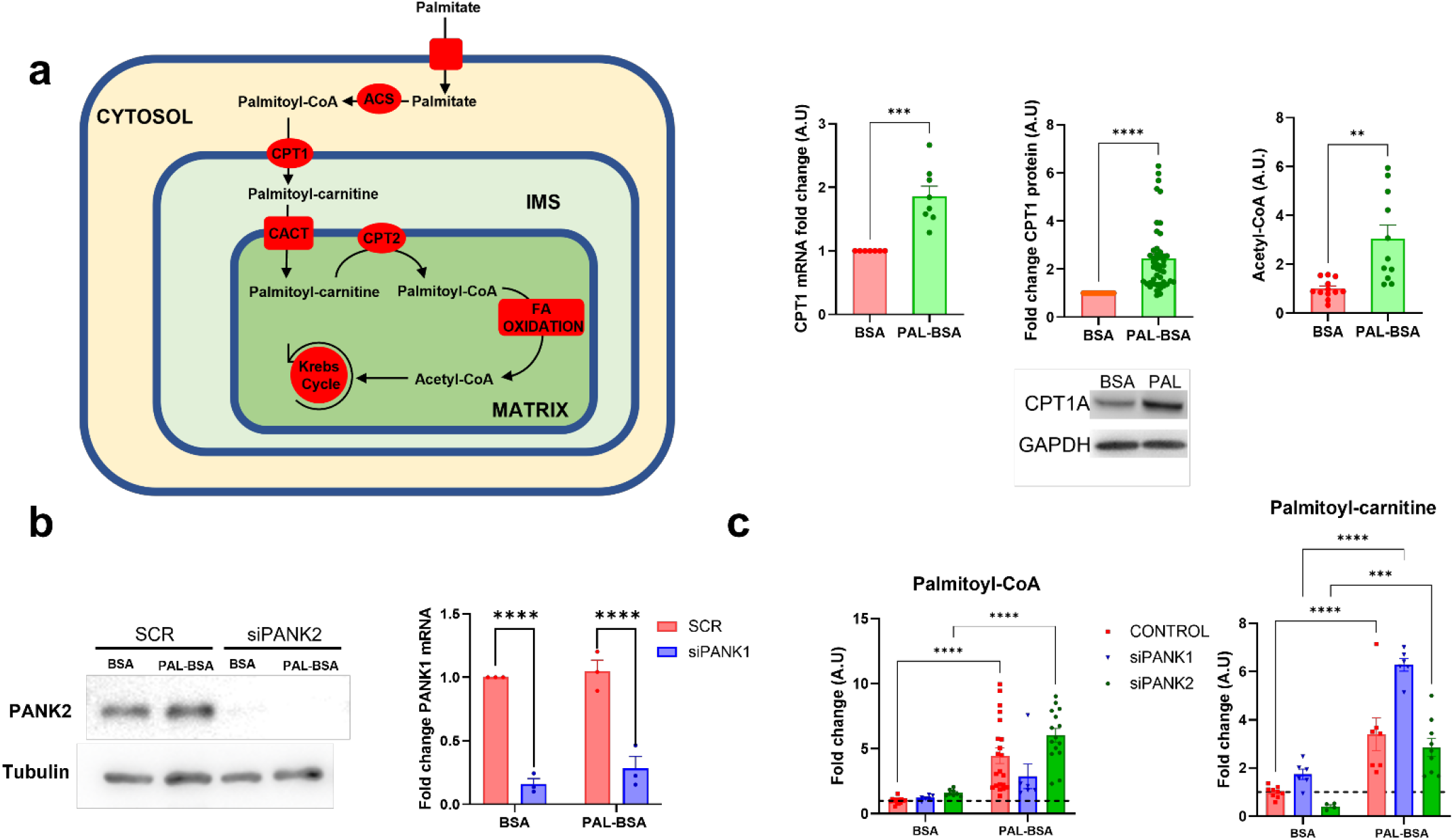
Exposure to PAL-BSA switch cellular metabolism towards Fatty Acid Oxidation. **a**, HEK293T cells were incubated with 250 μM BSA or palmitate-BSA (PAL-BSA) in DMEM + 10% FBS for 24 h. CPT1A mRNA and protein expression was measured by Quantitative PCR and western blot, respectively. Cellular levels of Acetyl-CoA measured by LC-MS. Incubation with PAL-BSA increased Acetyl-CoA production as expected. **b**, cells were incubated with Scramble (SCR) or specific siRNAs targeting PANK1 (SR325087BL) or PANK2 (SR325087BL). PANK2 expression was quantified by western blot and normalized to Tubulin expression. PANK1 expression was analyzed by measuring mRNA levels by Quantitative PCR. **c**, Total cellular levels of Palmitoyl-CoA and Palmitoyl-carnitine were analyzed by LC-MS. Mean ± SEM; n≥5 independent experiments. Significance assessed by 2-way ANOVA; *=p≤0.05, **=p≤0.01, ***=p≤0.001, ****=p≤0.0001. **ACS:** Acyl-CoA synthetase, **CPT1/2:** Carnitine Palmitoyltransferase 1 and 2, **CACT:** Carnitine Acyl-carnitine Translocase, **IMS:** Intermembrane Space.

**Figure 2.**
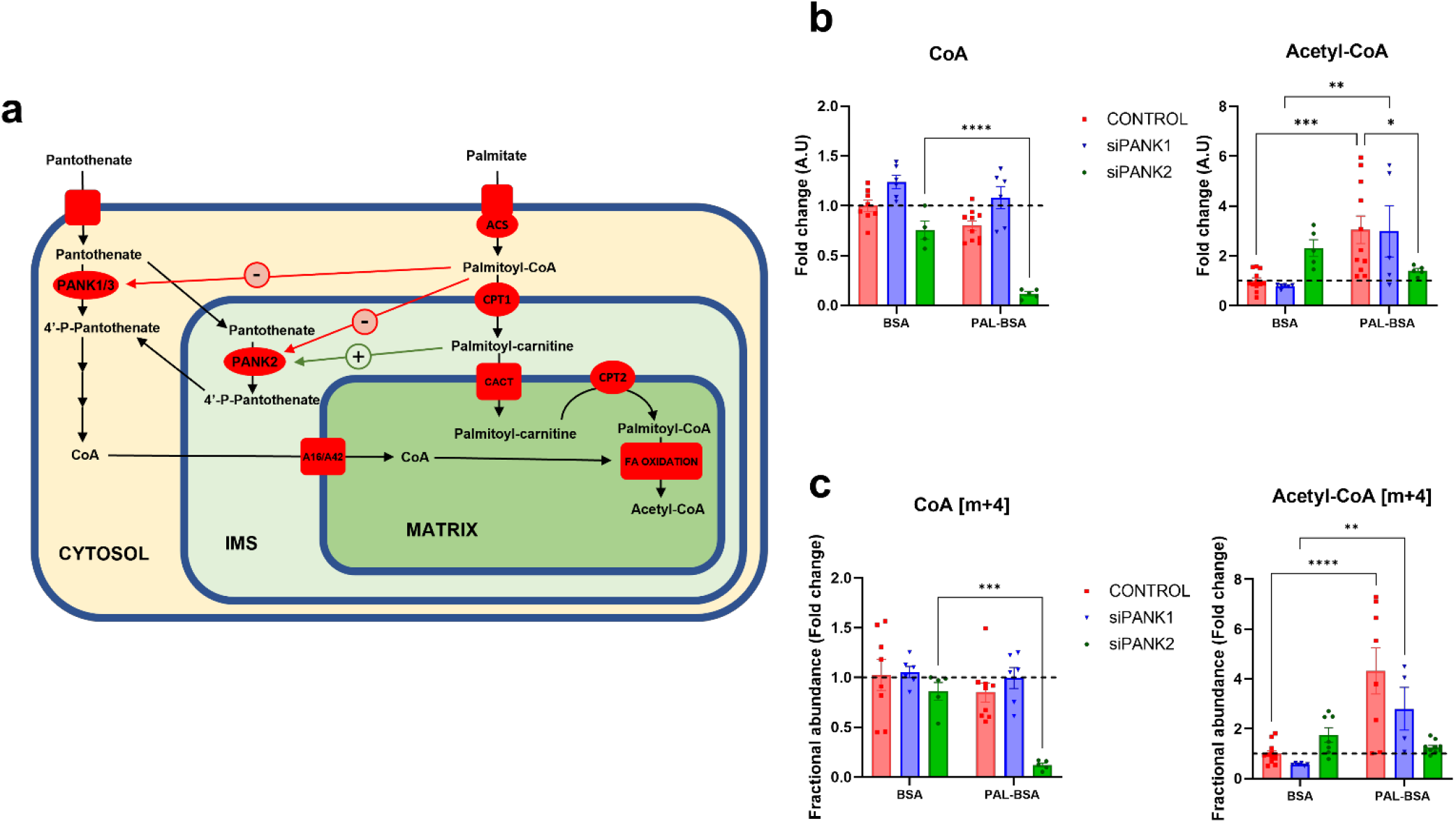
Diminished CoA synthesis in PANK2 deficient cells is uncovered by PAL-BSA incubation. **a**, Schematic representation of the role of PANK1-3 on CoA biosynthesis and the regulation of their activities. Control, siPANK1 or siPANK2 were incubated with 250 μM BSA or palmitate-BSA (PAL-BSA) in DMEM + 10% FBS for 24 h. 50 μM[^13^C_315_N_1_]-Pantothenate was pulsed for the last 6 hours of incubation. Cellular extracts were used to metabolite analysis by LC-MS. **b**, Total pool (unlabeled[M+0] + labeled [M+4]) of CoA and Acetyl-CoA. **c**, Fractional abundance (proportion of labeled (m+4)/total) of CoA and Acetyl-CoA. Total pools and fractional abundances of CoA decreased during PAL-BSA exposure only in PANK2 deficient cells, indicating diminished CoA synthesis in this condition. Incubation with PAL-BSA increased Acetyl-CoA production in Control and siPANK1 as expected. siPANK2 did not, suggesting an impairment of FAO, most likely due to diminished CoA synthesis in this condition. Mean ± SEM; n≥5 independent experiments. Significance assessed by 2-way ANOVA; *=p≤0.05, **=p≤0.01, ***=p≤0.001, ****=p≤0.0001. **PANK:** Pantothenase kinase, **ACS:** Acyl-CoA synthetase, **CPT1/2:** Carnitine Palmitoyltransferase 1 and 2, **CACT:** Carnitine Acyl-carnitine Translocase, **IMS:** Intermembrane Space. **A16/A42:** SLC25A16 and SLC25A42 mitochondrial carriers.

In order to analyze PANK-dependent CoA synthesis during PAL-BSA incubation, we performed tracing experiments using a stable isotope labeled pantothenate substrate in HEK293T control and PANK1 or PANK2-silenced cells (Figure 2). Cells were treated with scramble (CONTROL) or specific siRNAs targeting PANK1 (Origene, SR309983C) or PANK2 (Origene, SR325087B) and successful knockdown of *PANK1* or *PANK2* was confirmed by qPCR (*PANK1*) or western blot (PANK2) (Figure 1b). We cultured HEK293T controls, *PANK1* (siPANK1), or *PANK2* silenced (siPANK2) cells in the presence of vehicle BSA or BSA-palmitate (PAL-BSA) for 24 hours. Cultures were incubated with 50 μM [^13^C_3_,^15^N_1_]-pantothenate for the last 6 hours of treatment (Figure 2a). Interestingly, control and siPANK1 cells showed the same behavior. PAL-BSA treatment increased acetyl-CoA levels by approximately 3-fold in both controls and siPANK1 cells (Figure 2b), again consistent with an activation of fatty acid oxidation. No changes in CoA were evident in either controls or siPANK1 with PAL-BSA incubation (Figure 2b). However, incubation of siPANK2 cells with PAL-BSA led to a dramatic decrease in the cellular CoA pool (Figure 2b). Acetyl-CoA levels in *PANK2* knockdown cells were ∼2-fold higher than in controls under basal (BSA) conditions; however, PAL-BSA exposure did not trigger an increase in acetyl-CoA abundance in siPANK2 cells (Figure 2b).

Further analysis of m+4 isotopes showed that while the fractional abundance of [m+4] labeled CoA was stable in both controls and siPANK1 cells during either control conditions or PAL-BSA incubation, the fractional abundance of [m+4] acetyl-CoA increased concomitantly with total acetyl-CoA (Figure 2c). This suggests that under normal physiologic conditions, cellular *de novo* CoA synthesis from [^13^C_3_,^15^N_1_]-pantothenate should increase in order to keep up with metabolic demand during the activation of fatty acid oxidation. We found that siPANK2 cells failed to increase [m+4] labeled CoA (Figure 2c) despite increased demand, suggesting that *de novo* CoA synthesis is unable to keep up with demand. Consistently, siPANK2 cells were unable to increase [m+4] acetyl-CoA fractional abundance during PAL-BSA incubation (Figure 2c), indicating that FAO would be diminished in PANK2-deficient cells due to their limited capacity to synthesize new CoA needed to meet the demand to produce a larger pool of acetyl-CoA, with a critical role for PANK2 in maintaining CoA synthesis during metabolic switching to fatty acid oxidation.

Taken together these results indicate an important role for PANK2 in maintaining adequate CoA synthesis in the presence of high concentrations of long chain fatty acid carbon substrates. Its kinetic properties, specifically its stringent inhibition by acyl-CoAs that can be relieved by palmitoyl-carnitine (and possibly other long chain acyl-carnitines), place PANK2 in an optimal position to supply the appropriate amount of CoA based on the amount of substrate (fatty acids) to be oxidized (Figure 2a).

### PANK2 deficiency is associated with hallmarks of impaired FAO and starvation during PAL-BSA exposure

To investigate the metabolic effects of PANK2 deficiency we performed a targeted metabolic profiling of fatty acids by GC-MS. This analysis showed that siPANK2 cells accumulate both odd and even chain fatty acids upon PAL-BSA exposure compared to controls (Figure 3a). Interestingly, PANK1 silencing did not lead to fatty acid accumulation, indicating distinct roles for PANK1 and PANK2.

**Figure 3.**
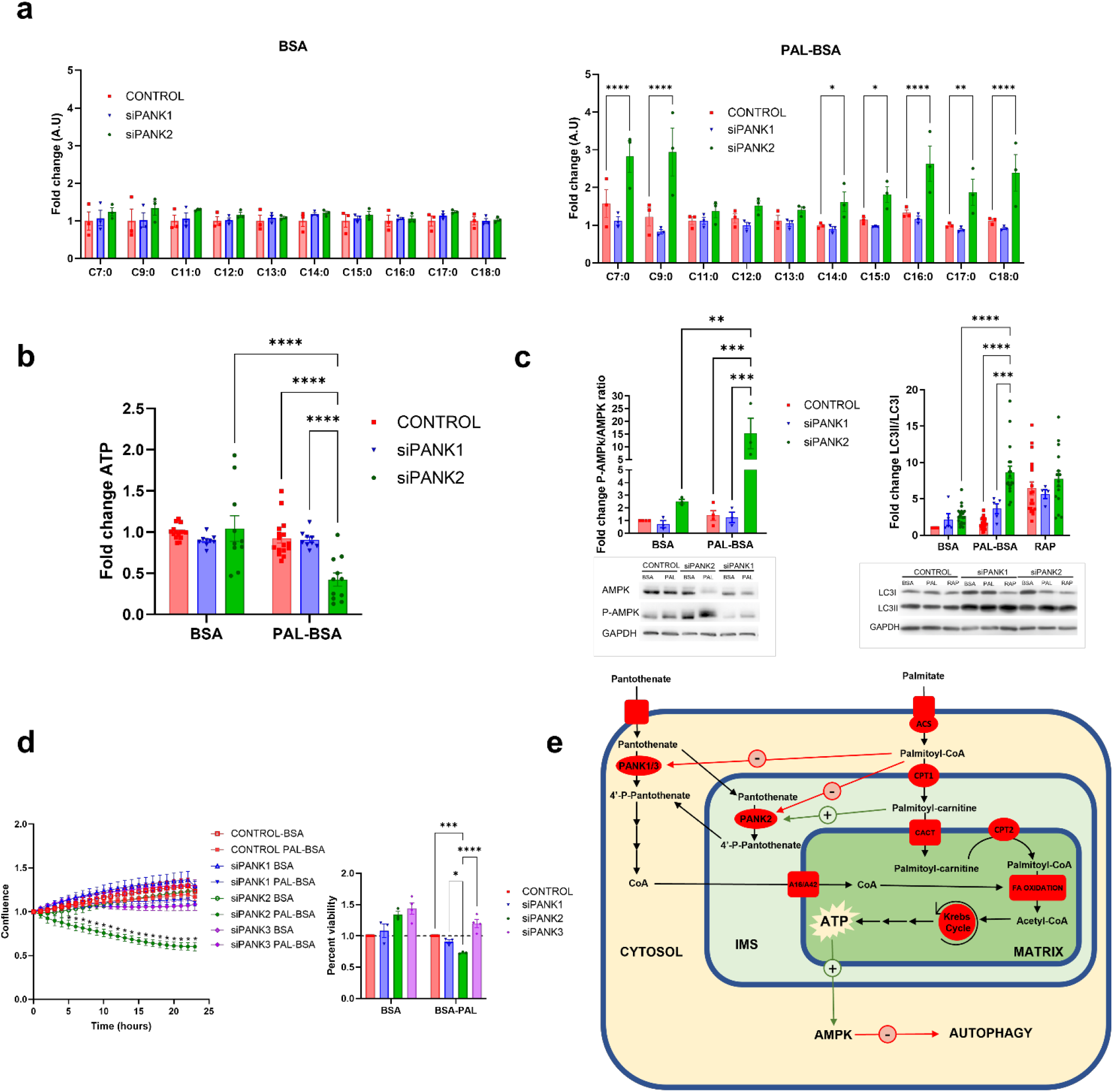
siPANK2 cells show metabolic impairment and starvation phenotypes during PAL-BSA treatment. HEK293T cells were treated with scramble (CONTROL) or specific siRNAs targeting PANK1 (SR325087BL), PANK2 (SR325087BL) or PANK3 (Origene, SR312489A). Cells were treated with either 250 μM BSA or PAL-BSA in DMEM + 10% FBS for 24h. Upon PAL-BSA exposure, siPANK2 cells showed hallmarks of FAO impairment, energy depletion and starvation. **a**, Analysis of Long Chain Free Fatty Acids by GC-MS. **b**, Cellular ATP levels. **c**, Analysis of the activation of AMPK pathway measured by the ratio of phosphorylated-AMPK/total AMPK., and Autophagic flux measured by the ratio of LC3II/LC3I. **d**, cell confluence measured with Incucyte and cell viability after 24 hours treatment analyzed by Trypan Blue exclusion assay. **e**, Schematic representation of the molecular pathways involved in CoA synthesis, energy generation from FAO. Mean ± SEM, n≥3 independent experiments. Significance assessed by 2-way ANOVA. *=p≤0.05, **=p≤0.01, ***=p≤0.001, ****=p≤0.0001. **PANK:** Pantothenase kinase, **ACS:** Acyl-CoA synthetase, **CPT1/2:** Carnitine Palmitoyltransferase 1 and 2, **CACT:** Carnitine Acyl-carnitine Translocase, **IMS:** Intermembrane Space. **A16/A42:** SLC25A16 and SLC25A42 mitochondrial carriers.

To better understand the cellular phenotypes associated with the loss of PANK2 and how it differs from the loss of PANK1, we next assessed ATP levels. We observed a decrease in ATP levels in PANK2-silenced cells (Figure 3b), consistent with impairment of fatty acid oxidation. The ATP reduction in siPANK2 was associated with an increase in bioenergetic stress master sensor AMP kinase activity (Figure 3c). Both siPANK1 and siPANK2 cells showed evidenced of increased autophagy activation measured by the ratio of LC3II/LC3I (Figure 3c) in both control and PAL-BSA conditions, however only siPANK2 cells dramatically increased the activation of autophagy (3.5-fold change) during PAL-BSA exposure compared to basal conditions. Taken together, these results indicate that siPANK2 cells struggle to maintain bioenergetic homeostasis during PAL-BSA exposure, leading to the activation of molecular pathways associated with cellular responses to starvation (Figure 3e). To test whether these cellular phenotypes influenced viability, we tracked cell confluence and cell viability in HEK293T cells transfected with PANK1 or PANK2 siRNA. In parallel, we silenced PANK3 (Supplemental Figure 1) to account for potential redundancy of function. Only in siPANK2 cells did PAL-BSA incubation diminish cell confluence and viability (Figure 3d); neither siPANK1 nor siPANK3 cells showed a significant decreased of confluence or viability, indicating that the PANKs are non-redundant and that the observed phenotype is specific to states of PANK2 deficiency. Moreover, we confirmed that this phenotype was consistently observed across multiple cell types, including lines of epithelial (HT1080) and neuronal (RenCell neural progenitor cell (NPC)) origin (Supplemental Figure 2).

Exposure to excessive fatty acids, particularly long chain saturated fatty acids, can lead to lipotoxicity, a condition characterized by excessive superoxide generation leading to endoplasmic reticulum stress and mitochondrial dysfunction [27-28]. It is known that high concentrations of palmitate trigger lipotoxicity via reactive oxygen species generation that leads to lipid peroxidation characterized by increased production of malondialdehyde (MDA) [29]. To test whether the observed phenotypes in siPANK2 are a result of lipotoxicity, control and siPANK2 cells were exposed to both palmitate-BSA and free palmitate (non-BSA conjugated). Tracking cell confluence revealed that free palmitate (PAL-NC) led to similar decreases in confluence in both control and siPANK2 cells, while only siPANK2 cells showed a significant decreased in confluence when exposed to PAL-BSA (Supplemental Figure 3). We measured MDA production in both control and siPANK2 cells and discovered that while exposure to PAL-BSA did not elicit any significant changes, free palmitate led to a dramatic production of MDA under both conditions (Supplemental Figure 3). This indicates that while free PAL exposure leads to nonspecific lipotoxicity in both controls and siPANK2 cells, the decreased confluence observed in siPANK2 cells does not appear to be related to MDA-associated lipotoxicity.

Finally, we wanted to test whether our findings could translate to PKAN models. Therefore, we used fibroblast lines derived from PKAN patients carrying a homozygous c.570C>A mutation that results in a premature stop codon at amino acid position 190 (p.Y190*). This mutation has been shown to lead to the total absence of PANK2 protein likely due to Nonsense-mediated mRNA decay [30]. Patients carrying this mutation usually present a classic, early-onset PKAN, which is severe and progressive [31]. Controls fibroblasts and the two PKAN lines were permeabilized and their capacity to oxidize palmitoyl-carnitine as a substrate for mitochondrial respiration was assessed using a Seahorse XFe24 (figure 4a). Results shown in Figure 4a demonstrated that PKAN-derived fibroblasts show impaired oxidative phosphorylation when palmitoyl-carnitine is used as a primary carbon source. Moreover, the same PKAN fibroblast lines showed a general capacity to oxidize the substrates included in regular DMEM medium (glucose, pyruvate and glutamine) similar to control cells (Figure 4b). These results extend our findings and indicate an impairment of FAO in multiple models of PANK2 deficiency.

**Figure 4.**
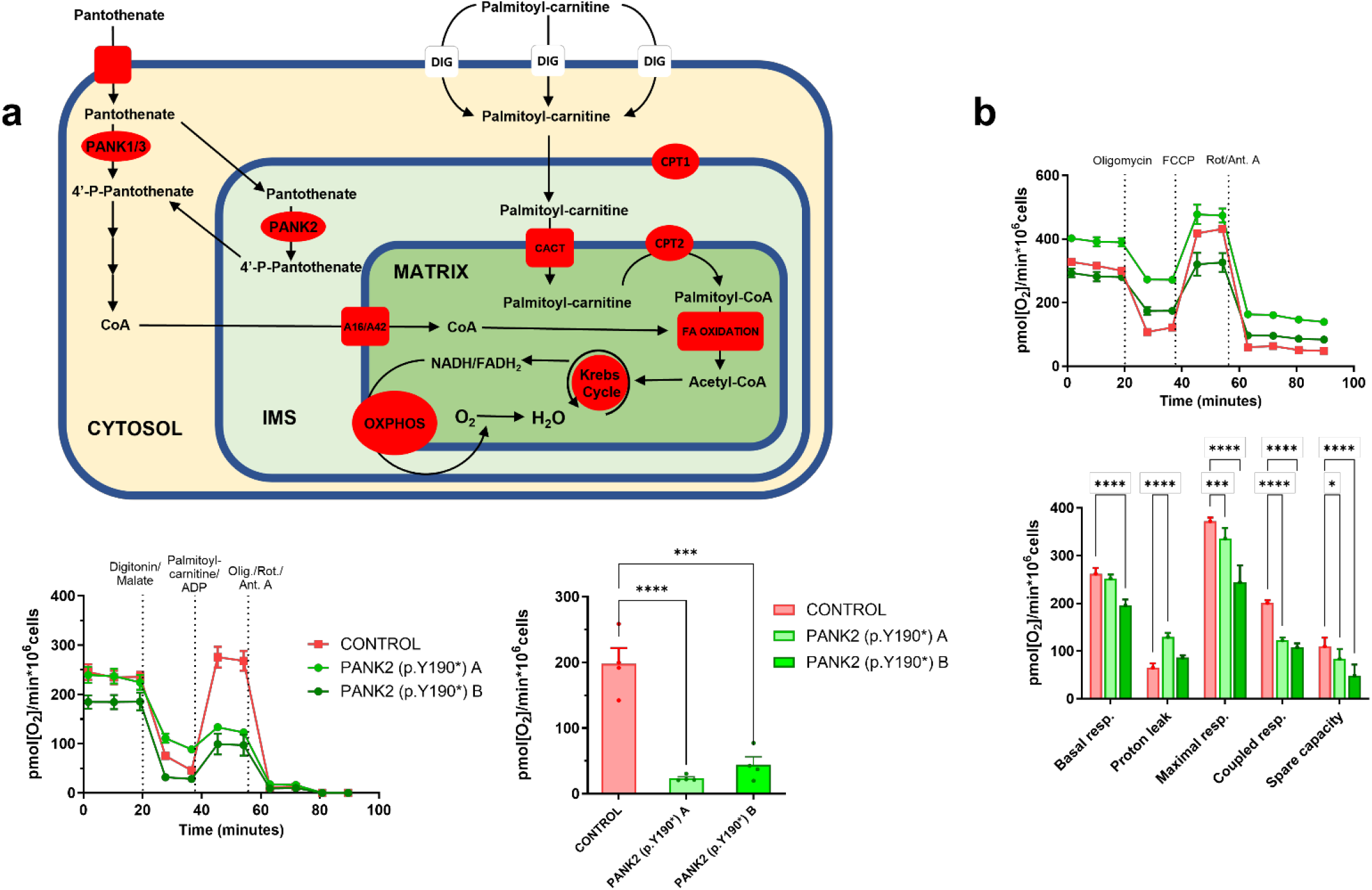
Fibroblasts from PKAN patients show impaired Fatty Acid oxidation. Control and PKAN derived fibroblasts were cultured in DMEM + 10% FBS. **a**, Fatty acid oxidation was analyzed by permeabilizing with 5 mg/ml digitonin + 250 mM malate and incubating with 50 mM palmitoyl-carnitine (bypassing CPT-I) and 1 mM ADP. **b**, Mitochondrial respiration using Glucose as substrate. Mitochondrial Inhibitors 1 mM oligomycin, 1 mM rotenone, and 2.5 mM antimycin A were added as indicated. OCR measured by SeaHorse XF24. Mean ± SEM, n≥3 independent experiments. Significance assessed by 2-way ANOVA. *=p≤0.05, **=p≤0.01, ***=p≤0.001, ****=p≤0.0001. **PANK:** Pantothenase kinase, **CPT1/2:** Carnitine Palmitoyltransferase 1 and 2, **CACT:** Carnitine Acyl-carnitine Translocase, **IMS:** Intermembrane Space. **A16/A42:** SLC25A16 and SLC25A42 mitochondrial carriers, **OXPHOS:** Oxidative Phosphorylation

Collectively, our findings demonstrate that a distinct and essential role for PANK2 in *de novo* CoA synthesis during conditions of metabolic switching to cellular energy production dependent on fatty acid oxidation.

## DISCUSSION

Our results highlight the essential nature of PANK2’s function when lipid catabolism is activated. We present evidence that this role is specific to PANK2 among the PANKs, as knocking down either PANK1 or PANK3 did not lead to diminished viability (Figure 3 and Supplemental Figure 2) during PAL-BSA incubation. Previous cell-free *in vitro* studies have already hinted at this, showing that PANK2 feedback inhibition by acetyl-CoA and palmitoyl-CoA can be relieved by binding palmitoyl-carnitine, which increases when fatty acid oxidation is activated, especially in the mitochondrial intermembrane space (IMS) where the PANK2 protein resides [17].

We show in figure 1 that incubation with BSA-conjugated palmitic acid (PAL-BSA) establishes the appropriate cellular conditions for PANK2 activation. Moreover, results shown in Figure 2 demonstrate that PANK2, but not PANK1, is essential during this condition to maintain an appropriate CoA synthesis that meets the demand for increased acetyl-CoA production through the oxidation of fatty acids. These results together with previous *in vitro* and protein localization studies indicate that PANK2 is the main driver of CoA synthesis during fatty acid oxidation because: 1) PANK2 possess the optimal biochemical properties (the ability to reverse acyl-CoA-mediated feedback inhibition by binding palmitoyl-carnitine) to maintain its enzymatic activity in a cellular environment where palmitoyl-CoA levels are high [17], and 2) PANK2 localization within the mitochondrial IMS [16] where local acyl-carnitine concentrations are high due to the activity of CPT1A given its role in transporting fatty acids to the mitochondrial matrix to undergo oxidation. In this working model (Figure 2a), a lack of PANK2 activity during conditions where fatty acid oxidation predominates could result in the impairment of this process and failure to maintain appropriate cellular bioenergetics (Figure 3e). Supporting this hypothesis, PANK2 silenced cells, but not siPANK1 or siPANK3, showed fatty acid accumulation, decreased cellular viability, low ATP alongside activation of AMPK and autophagy during PAL-BSA incubation, consistent with a cellular starvation phenotype and impaired bioenergetic status despite abundant nutrients (Figure 3).

A recent review by Hayflick, et al. questioned the widespread assumption that PKAN is caused by depletion of CoA, as no cellular CoA deficiency has been found in cells derived from human PKAN patients, in PANK2 knockout mice, or in human cell lines with knockdown of PANK2, including neurons [19]. Our studies have uncovered cellular conditions that cause a disruption of *de novo* CoA synthesis and lead to cellular CoA deficiency in a PANK2-deficient model. Previous studies have shown higher expression of PANK1 and PANK3 in PKAN fibroblasts compared to controls [32-33], leading to the assumption that this is the main reason PKAN models do not show a deficiency in CoA. We found an increase in both *PANK1* and *PANK3* expression in siPANK2 cells (Supplemental Figure 4), yet despite this adaptation, PAL-BSA incubation led to a CoA deficiency in siPANK2 indicating that solely increasing the expression of other PANKs is not enough to restore CoA levels after the loss of PANK2. Interestingly, a recent study has brought new insights to the field by discovering that PANK4 is a phosphatase that actively removes the CoA synthesis intermediate 4-phosphopantethine and effectively reduces the production of the final product, indicating that cellular CoA content is tightly controlled [18]. Moreover, the authors demonstrated that the insulin-PI3K-Akt axis, a major regulator of metabolism and growth, stimulates *de novo* CoA synthesis by phosphorylating PANK4 and inhibiting its phosphatase activity, solidifying a previously suggested link between PI3K signaling and CoA synthesis [34] and suggesting that other additional adaptations may be needed in order to restore cellular CoA levels after PANK2 loss.

A recent study has confirmed that CoA biosynthesis is a process that primarily occurs in the cytosol while the mitochondrial CoA pool is established by the action of SLC25A16 and SLC25A42 mitochondrial carriers [35]. Such a mitochondrial CoA pool is critical for supporting intra-mitochondrial FAO. Data presented in the current work complement the previous study suggesting that PANK2 might be especially important to maintain the mitochondrial CoA pool and indicating that the combined action of PANK2 and CoA mitochondrial carriers is needed to effectively meet the high demand of mitochondrial CoA that FAO requires. Notably, the homozygous N291D mutation in SLC25A42 has been shown to cause mitochondrial encephalomyopathy [36-37].

The experiments presented in this paper demonstrate for the first time the unique cellular role of PANK2 in CoA biosynthesis. Our results highlight the importance of PANK2 activity when lipid catabolism is activated. Moreover, we found evidence of impaired FAO in PKAN-derived fibroblasts supporting our findings across multiple cell types and suggesting the same metabolic abnormalities may be present in PKAN patients. Understanding the distinct roles of the PANKs as regulators of cellular metabolism may have important implications for cancer, cardiovascular disease, and neurodegeneration

## METHODS

### Cell culture

HEK293T, HT1080 and Human Fibroblast cell lines were maintained in complete growth media containing DMEM (Life Technologies) supplemented with 10% FBS and 1mM GlutaMAX (Gibco), 1mM sodium pyruvate (Sigma) and 1% penicillin/streptomycin at 37°C under 5% carbon dioxide with a humidified atmosphere. ReNcell culture: ReNcell human NPCs were maintained in an undifferentiated state by culturing in ReNcell maintenance medium supplemented with 1X B-27, 25 ng/mL EGF, 25ng/mL bFGF and 1% penicillin/streptomycin, on Matrigel-coated T-25 flasks.

### siRNA knockdown

Cells were seeded at a density of 0.3 × 10^6^ cells per 6 well dish the day before transfection. Two transfections were performed on sequential days using fresh OptiMEM (Gibco) supplemented with 6.5% FBS. Transfections followed the Lipofectamine RNAiMAX (ThermoFisher) protocol with control scramble siRNA (Origene, SR30004), or human siRNA oligo duplex against either PANK2 (Origene, SR325087B, Sequence: 5’GUAUUUUUCUAAGUCAUCAAGAUAA3’), PANK1 (Origene, SR309983C, sequence: 5’AAUGACUGAUGACAAGUAGAGACGA3’) or PANK3 (Origene, SR312489A, sequence: 5’CAAGAGCUACUUUAGUUACUAUCAC3’). On the fourth day, cells were provided with fresh complete growth media supplemented with either vehicle BSA or palmitic acid conjugated to BSA at a 250µM final concentration. Cells were harvested after the 24 hour PAL-BSA exposure.

### Autophagic flux measurements

Autophagy was assessed by tracking the conversion of LC3-I to LC3-II, with and without the lysosomal inhibitor chloroquine (CQ) (Sigma) at 100nM final concentration for 4 hours. Rapamycin was used at 200nM in some experiments to induce autophagy.

### RIPA buffer protein extraction and western blotting

Cellular proteins were extracted with RIPA buffer (Thermo Fisher) supplemented with protease (Thermo Fisher) and phosphatase inhibitor cocktails (Sigma) on ice and centrifuged at 13,000g. The supernatant was collected. Protein concentrations were measured with the BCA assay and an equal amount of protein separated on SDS-page and transferred to a 0.2µm nitrocellulose membrane (BIO-RAD). Membranes were blocked with 5% BSA. The membranes were incubated with antibodies against anti-LC3B (Abcam; ab51520) at 1:5000 dilution, anti-AMPK (Cell signaling; 2532) at 1:1000 dilution, anti-phospho-AMPK (Cell signaling; 2531) at 1:5000 dilution, anti-CPT1A (Abcam; ab128568) at 1:10000 dilution, anti-PANK2 (Origene; TA501419) at 1:2000 dilution, anti-tubulin (Abcam; ab4074) at 1:10000 dilution, or anti-GAPDH (Abcam; ab9485) at 1:5000 dilution and detected using anti-rabbit or anti-mouse HRP (GE Health Sciences) at a 1:5,000 dilution.

### Cell counts and confluence measurements

Growth of siRNA-treated cells with either vehicle BSA or PAL-BSA were monitored with the Incucyte S3 (Sartorius) at 10X and analyzed with the confluence setting for 24 hours. Percent live cells and cell numbers were calculated based on trypan blue staining and counted with a Countess II FL cell counter (Life Technologies). For the cell viability experiment presented in Supplementary Figure 2, LIVE/DEAD ™ viability/cytotoxicity kit from Thermofisher (Cat # L3224) was used according to the manufacturer’s instructions. 2-D live cell imaging was performed using the Keyence BZ-X800E with a 40x objective. Cells were analyzed and counted manually using ImageJ.

### qPCR assays

RNA was extracted using a Direct-zol RNA MiniPrep kit (Zymo Research) according to the manufacturer’s instructions, with DNase I (Zymo Research) treatment. Reverse transcription was done with iScript™ Reverse Transcription Supermix for RT-qPCR (BIO-RAD) following the manufacturers protocol. qPCR was performed with SsoAdvanced Universal SYBR Green Supermix (BIO-RAD) following the manufacturer’s protocol. Assay IDs were GAPDH Hs.PT.39a.22214836, PANK1 Hs.PT.58.2284965, PANK3 Hs.PT.58.2201075, all from IDT.

### ATP quantification

ATP from 10^6^ cells was quantified via luminescence using an ATP quantification kit (Abcam; ab113849).

### Lipid peroxidation (MDA) quantification

MDA from 10^6^ cells was quantified using a commercial kit (Abcam; ab233471) following manufacture’s specifications.

### Oxygen consumption measurements with Seahorse 24XFe analyzer

Oxygen consumption rate (OCR) was measured with the Seahorse extracellular flux 24XFe analyzer (Agilent). Cells were seeded at 50,000 cells/well on 24-well culture plates (Agilent) 24-hours prior to measurements. An hour before measurements the media was replaced with Seahorse XF media supplemented with 1mM glucose, 1mM glutamine, and 1mM sodium pyruvate incubated at 37 without CO_2_. Baseline OCR was measured before injecting inhibitory compounds and then followed by OCR measurements after the sequential addition of oligomycin (1µM), FCCP (2µM), and antimycin (2.5µM)/rotenone (1µM) respectively. The OCR measurements were normalized to cell numbers counted with the Scepter™ 2.0 cell counter.

#### Palmitoyl-carnitine oxygen consumption measurements

Cells were seeded at 50,000 cells/well on 24-well culture plates (Agilent) 24-hours prior to measurements. An hour before measurements the media was replaced with a mitochondrial assay solution (MAS) containing HEPES, MgCl_2_, KH_2_PO_4_, mannitol, sucrose and EGTA (pH 7.2) and BSA was added fresh the same day. The first injection contained digitonin (5µg/ul) and malate (250mM). Next, injection contained the palmitoyl-carnitine (50µM) and ADP (1mM). Finally, antimycin A (2.5µM) and rotenone (1µM) was injected to inhibit mitochondrial respiration. The OCR measurements were normalized to cell numbers counted with the Scepter™ 2.0 cell counter.

### CoA, Acetyl-CoA, and fatty acid measurements

Cells were grown in 6 well plates with confluence tracked with using the Incucyte for normalization for 24 hours, then washed with ice cold PBS followed by H_2_O. The plates were snap frozen in liquid nitrogen for GC-MS (fatty acid analysis) and LC-MS (CoA and carnitine species) measurements. Cell culture plates were lyophilized overnight and then scraped into 1 ml of ice-cold 2:2:1 methanol:acetonitrile:water containing a mixture of internal standards (D4-citric acid, D4-succinic acid, D8-valine, and U13C-labeled glutamine, glutamate, lysine, methionine, serine, and tryptophan; Cambridge Isotope Laboratories) to extract metabolites. The cell extraction mixtures were transferred to a microcentrifuge tube and flash frozen in liquid nitrogen. Frozen extracts were thawed for 10 minutes in a water bath sonicator and then incubated for 1 hour at – 20°C. Metabolite extracts were centrifuged for 10 minutes at 21,000 x g and supernatants were transferred to autosampler vials (GC) or microcentrifuge tubes (LC) and dried using a SpeedVac vacuum concentrator (Thermo).

#### GC-derivatization

Dried metabolite extracts were derivatized using methoxyamine hydrochloride (MOX) and N,O-Bis(trimethylsilyl)trifluoroacetamide (TMS). Briefly, dried extracts were reconstituted in 30 μl of 11.4 mg/ml MOX in anhydrous pyridine, vortexed for 5 minutes, and heated for 1 hour at 60°C. Next, 20 μl of TMS was added to each sample, samples were vortexed for 1 minute, and heated for 30 minutes at 60°C.

#### GC-MS method for fatty acid quantification

Derivatized samples were analyzed using GC-MS. GC was conducted using a Trace 1300 GC (Thermo) fitted with a TraceGold TG-5SilMS column (Thermo). 1 μl of derivatized sample was injected into the GC operating under the following conditions: split ratio = 20-1, split flow = 24 μl/minute, purge flow = 5 ml/minute, carrier mode = Constant Flow, and carrier flow rate = 1.2 ml/minute. The GC oven temperature gradient was as follows: 80°C for 3 minutes, increasing at a rate of 20°C/minute to 280°C, and holding at a temperature at 280°C for 8 minutes. Ion detection was performed by an ISQ 7000 mass spectrometer (Thermo) operated from 3.90 to 21.00 minutes in EI mode (−70eV) using select ion monitoring (SIM).

#### LC-sample preparation for CoA and acetyl-CoA measurements

Dried metabolite extracts were reconstituted in a 1/10th volume of 1:1 acetonitrile:water and vortexed for 10 minutes (200 μl of metabolite extract was resuspended in 20 μl). Reconstituted extracts were maintained at −20°C overnight and centrifuged at 21,000 x g for 10 minutes. Supernatants were transferred to autosampler vials for LC-MS analysis.

#### LC-MS method

2 µL of the prepared samples were separated using a Millipore SeQuant ZIC-pHILIC (2.1 × 150 mm, 5 µm particle size) column with a ZIC-pHILIC guard column (20 x 2.1 mm) attached to a Thermo Vanquish Flex UHPLC. Mobile phase was comprised of Buffer A [20 mM (NH4)_2_CO_3_, 0.1% NH_4_OH (v/v)] and Buffer B [acetonitrile]. The chromatographic gradient was run at a flow rate of 0.150 mL/min as follows: 0–21 min-linear gradient from 80 to 20% Buffer B; 20-20.5 min-linear gradient from 20 to 80% Buffer B; and 20.5–28 min-hold at 80% Buffer B.

#### High Resolution Mass Spectrometry

The mass spectrometer was operated in full-scan, polarity-switching mode from 1 to 20 minutes, with the spray voltage set to 3.0 kV, the heated capillary held at 275 °C, and the HESI probe held at 350 °C. The sheath gas flow was set to 40 units, the auxiliary gas flow was set to 15 units, and the sweep gas flow was set to 1 unit. MS data acquisition was performed in a range of m/z 70–1,000, with the resolution set at 70,000, the AGC target at 1 × 10_6_, and the maximum injection time at 200 ms.

#### Data analysis

Raw data were analyzed using TraceFinder 4.1 (Thermo). For GC-MS data, metabolite identification and annotation required at least two ions (target + confirming) and a unique retention time per metabolite that corresponded to the ions and retention times of reference standards previously determined in-house. For LC-MS data, metabolite identification and annotation required ion accurate mass ± 5 mmu and retention time per metabolite to correspond to the ions and retentions times of reference standards previously determined in-house. A pooled-sample generated from metabolite extracts was analyzed before, at a set interval, and after the analytical run to correct peak intensities using the NOREVA tool [22].

### Isotope tracing analysis

For the analysis of isotopes of CoA and Acetyl-CoA after [^13^C_3_,^15^N_1_]-pantothenate incubation, LC sample preparation LC separation methods were the same as in the previous section. Injection volume was 8 μl.

#### High Resolution Mass Spectrometry

The mass spectrometer was operated in tSIM (targeted selected ion monitoring) mode from 1 to 20 minutes and directed against an inclusion list comprising ion-retention time couples generated from the University of Iowa Metabolomics Core facility standard-confirmed, in-house library with an isolation width of 12 m/z and an offset of 5 m/z. tSIM scan filters were multiplexed up to 8 times in negative polarity with the spray voltage set to 3.0 kV, the heated capillary was held at 275 °C, and the HESI probe was held at 350 °C. The sheath gas flow was set to 40 units, the auxiliary gas flow was set to 15 units, and the sweep gas flow was set to 1 unit. MS data acquisition was performed in a range of m/z 72–1,080, with the resolution set at 70,000, the AGC target at 5e4, and the maximum injection time at 200 ms [23].

#### Data analysis

Acquired LC-MS data were processed by Thermo Scientific TraceFinder 5.2 software, and metabolites were identified based on the University of Iowa Metabolomics Core facility standard-confirmed, in-house library. NOREVA was used for signal drift correction [22]. Data were normalized to the sum of all the measured metabolite ions in that sample

### Statistical analysis

Experiments were run at least in triplicate and repeated a minimum of three times using independent biological replicates. Student’s t-test was used when comparing the means of two groups, and one-/two-way ANOVA with Tukey’s post hoc when comparing the means of more than two groups. A limited number of statistical outliers were excluded after identification by Grubbs test. N values within figure legends represent independent experiments.

## Acknowledgements

We thank the University of Iowa Metabolomics Core Facility, Fraternal Order of Eagles Diabetes Research Center for their assistance with metabolic profiling and isotope tracing experiments

We thank Ana Sanchez Cuesta for her technical assistance with OCR analysis of human Fibroblasts.

MCK is supported in part by NIH R01 NS127108

## Contributions

S.M.N. S.P.L and M.C.K. conceived the study. S.M.N designed and performed experiments. S.P.L supervised the study, performed experiments and wrote the manuscript with input from all authors. R.S, A.M.S.E, B.A.W, U.H. and F.N provided technical support for the experiments performed in Kruer and Padilla-Lopez lab. E.B.T, A.R. and I.A.W. performed and helped with mass spectrometry analysis. C.S.O and A.P.R performed the experiments with PKAN fibroblasts.

## Data availability

Original data is available from the corresponding authors upon reasonable request.

## Supporting information

This article contains supplementary supporting information.

