## Supplemental figures for "PANK2-mediated *de novo* CoA synthesis is required for metabolic switching to fatty acid oxidation"

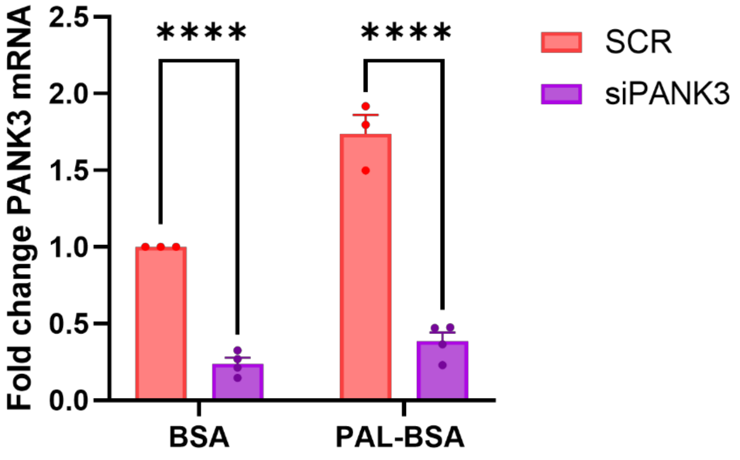


**Supp Figure 1. PANK3 siRNA treatment successfully reduced expression**. HEK293T cells were incubated with Scramble (SCR) or specific siRNAs targeting PANK3 (SR312489A). PANK3 expression was analyzed by measuring mRNA levels by Quantitative PCR. Cells were treated with either 250 µM BSA or PAL-BSA in DMEM + 10% FBS. Mean ± SEM, n≥3 independent experiments. Significance assessed by 2-way ANOVA. *=p≤0.05, **=p≤0.01, ***=p≤0.001, ****=p≤0.0001

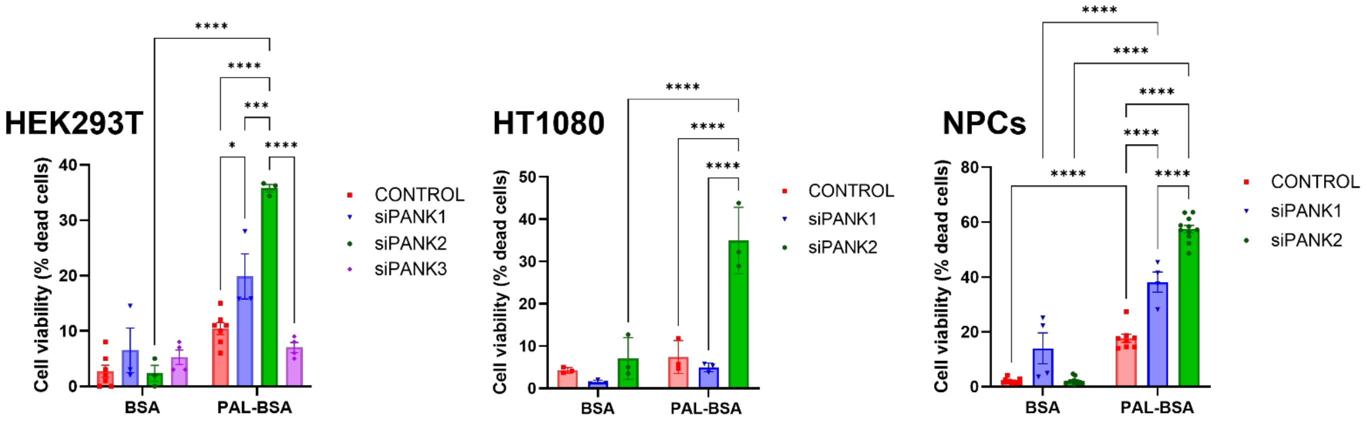


**Supp Figure 2. PAL-BSA treatment triggered viability loss in siPANK2 cells distinct from siPANK1 and siPANK3**. PANK1-3 were individually silenced in HEK293T cells with scramble (CONTROL) or specific siRNAs targeting PANK1 (Origene, SR325087BL), PANK2 (Origene, SR325087BL) and PANK3 (Origene, SR312489A). Cells were treated with either 250 μM BSA or PAL-BSA in DMEM + 10% FBS. Upon PAL-BSA exposure, siPANK2 cells showed increased cell death after 24 hours treatment analyzed by Tripan Blue exclusion assay.. Mean ± SEM, n≥3 independent experiments. Significance assessed by 2-way ANOVA. *=p≤0.05, **=p≤0.01, ***=p≤0.001, ****=p≤0.0001.

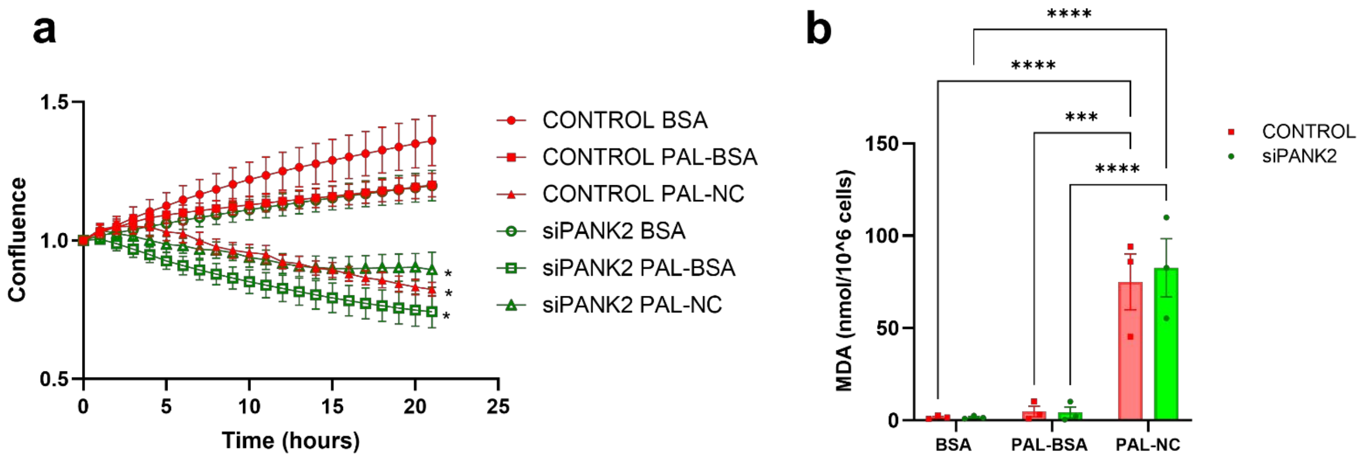


**Supp Figure 3. Free Palmitate treatment triggered cell confluence loss and MDA accumulation HEK293T cells**. HEK293T cells were treated with scramble (CONTROL) or specific siRNA targeting PANK2 (Origene, SR325087BL) Cells were treated with either 250 μM BSA, PAL-BSA or unconjugated Palmitate (PAL-NC) in DMEM + 10% FBS. PAL-NC led to a decreased of both cell confluence measured with Incucyte **(a)** and MDA production **(b)** in both control and siPANK2 cells. Mean ± SEM, n≥3 independent experiments. Significance assessed by 2-way ANOVA. *=p≤0.05, **=p≤0.01, ***=p≤0.001, ****=p≤0.0001.

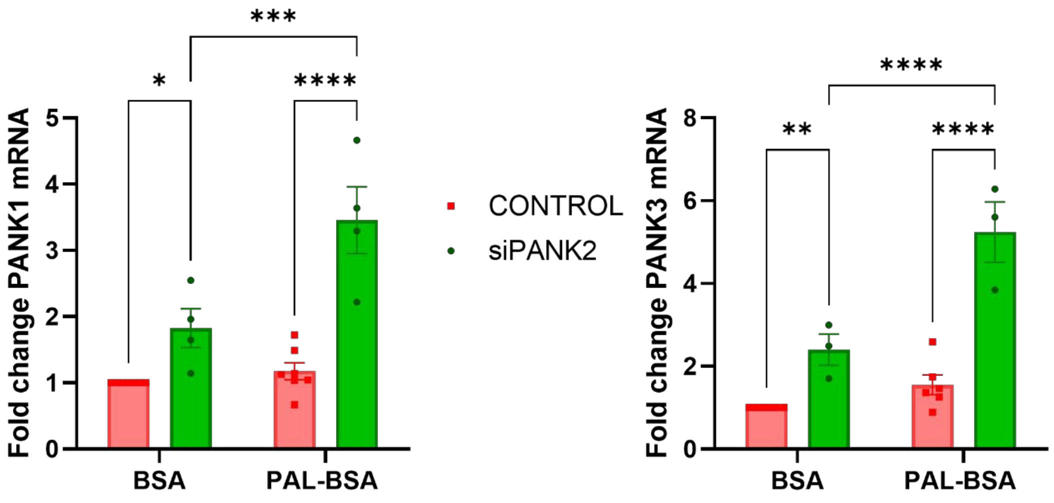


**Supp Figure 4. PANK1 and 3 expression increases in siPANK2 cells**. HEK293T cells were incubated with scramble (CONTROL) or specific siRNAs targeting PANK2 (Origene, SR325087B). Cells were treated with either 250 μM BSA or PAL-BSA in DMEM + 10% FBS. After treatment, PANK1 and PANK3 mRNA expression was analyzed by Quantitative PCR. Mean ± SEM, n≥3 independent experiments. Significance assessed by 2-way ANOVA. *=p≤0.05, **=p≤0.01,***=p≤0.001, ****=p≤0.0001.
